# Talker head-orientation and extended high-frequency benefits for speech recognition as a function of masker head angle

**DOI:** 10.64898/2026.08.12.744468

**Authors:** Vahid Delaram, Rohit M. Ananthanarayana, Allison Trine, Margaret K. Miller, G. Christopher Stecker, Emily Buss, Brian B. Monson

## Abstract

Several types of cues contribute to speech recognition in multi-talker environments. In this study, we investigated how talker head-orientation related (THOR) cues and extended high- frequency (EHF; >8kHz) cues affect speech-in-speech recognition for both female and male speech. We examined the THOR benefit associated with a non-facing masker talker head orientation (relative to a facing orientation) as a function of masker talker facing angle. The target talker always faced the listener, whereas co-located maskers were tested with eight different masker head angles, ranging from 0° (facing the listener) to facing 180° away. Two filtering conditions were tested: full- band and low-pass filtered at 8 kHz. A THOR benefit was observed at masker head angles greater than 45°, increasing from 2 dB to 8 dB between angles of 67.5° and 180°. This benefit was reduced for low-pass filtered speech. Access to EHF cues improved performance, but only for masker head angles >22.5°. There was no significant relationship between 16-kHz pure-tone thresholds and performance for young, normal-hearing listeners with good EHF hearing. These findings indicate that listeners benefit from non-facing masker talker head orientations >45° when the target talker is facing the listener, with greater benefit for larger head angles.

## I. INTRODUCTION

Multi-talker environments can pose a significant challenge for speech recognition as multiple speech signals compete for listener attention. The ability to accurately segregate and recognize speech in these complex environments, often referred to as the *cocktail party problem*, is a fundamental aspect of auditory scene analysis (Bronkhorst, 2000; Cherry, 1953). Several well-known factors influence this ability, including talker spatial position on the horizontal plane (Dirks and Wilson, 1969; Noble and Perrett, 2002), talker gender (Brungart et al., 2001; Darwin et al., 2003), and the number of talkers (Freyman et al., 2004; Rosen et al., 2013). Another lesser-known but important factor that influences speech recognition in multi-talker scenes is talker head orientation (Braza et al., 2022; Flaherty et al., 2021; Monson et al., 2019; Strelcyk et al., 2014).

Speech emanating from the mouth of a talker exhibits frequency-dependent radiation patterns. Lower-frequency components tend to propagate more omnidirectionally, whereas higher-frequency components tend to propagate more directionally, typically toward the front of the talker (Chu and Warnock, 2002; Dunn and Farnsworth, 1939; Leishman et al., 2021; Monson et al., 2012a; Monson and Ananthanarayana, 2023; Trine et al., 2025). Because of this speech directivity, the physical orientation of talkers’ heads relative to the listener determines the distribution of speech spectral energy delivered to the listener, creating talker head orientation-related (THOR) cues for talker head orientation discrimination (Moriarty et al., 2024). Most prior research evaluating speech-in-speech recognition has used stimuli recorded from a microphone directly in front of the talker, simulating a condition in which all talkers are facing the listener (Figure 1A). However, in everyday conversations, it would be unusual for all active talkers to be facing the listener. More commonly, the target talker faces the listener, while maskers (i.e., background talkers) face away from the listener, orienting toward their own conversation partners (Vertegaal et al., 2001; Figure 1B and 1C).

**Figure 1.**
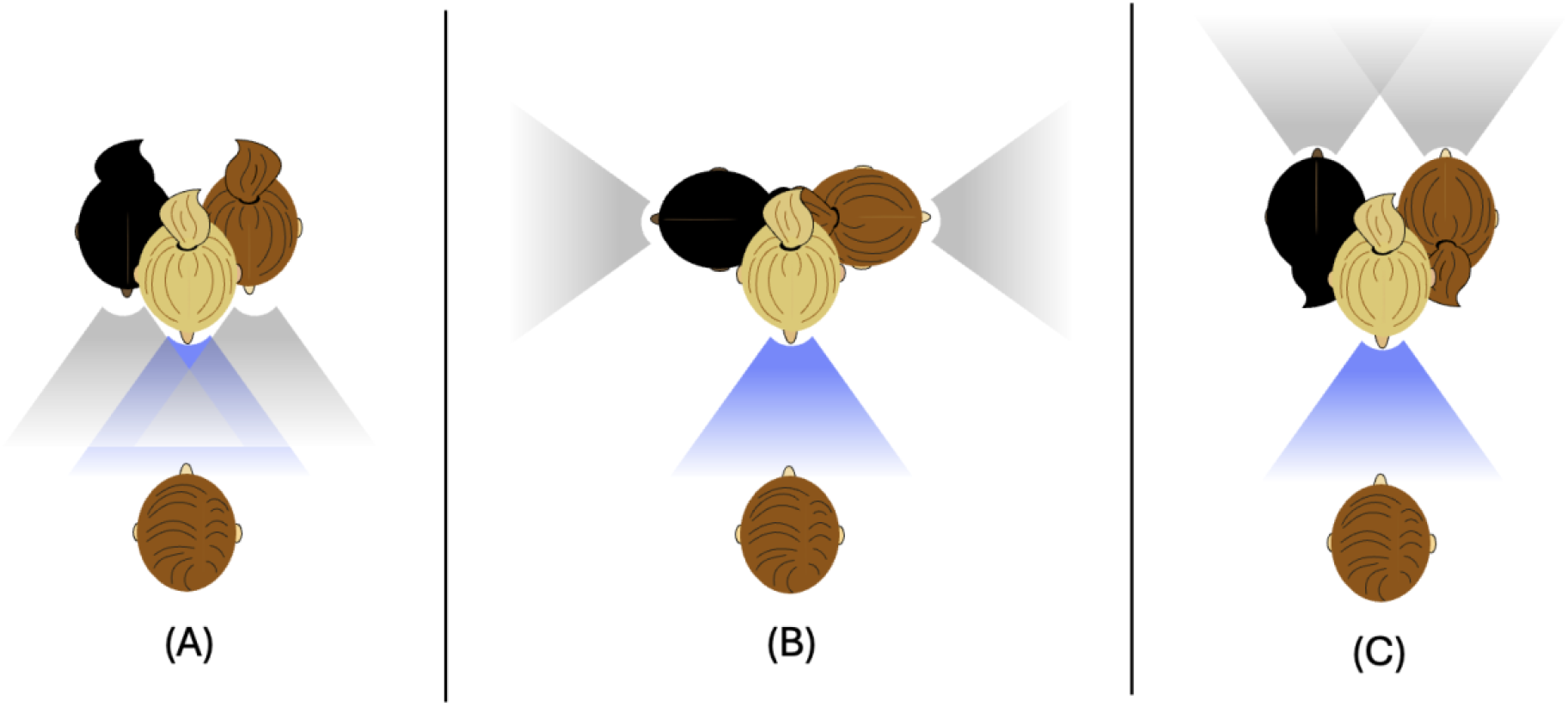
(A) All talkers are facing the listener. (B, C) The target talker is facing the listener, and the maskers are facing ±90° (B) or 180° away (C). Shaded beams depict talkers’ speech high-frequency energy.

THOR cues associated with non-facing maskers can influence speech recognition performance. Several studies have demonstrated improved speech recognition when the target talker is facing the listener and maskers face away from the listener. For example, Strelcyk et al. (2014) reported better recognition when spatially separated maskers were oriented 105° away from the listener compared to facing the listener (0°). Braza et al. (2022) also observed THOR benefits when changing masker head orientation angles from 0° to 60°. The largest THOR benefit occurred when all talkers were co-located on the horizontal plane, followed by symmetrical and asymmetrical spatial separation conditions. Ananthanarayana et al., (2024) demonstrated a THOR benefit comparing co-located masker head angles of 0° and 56.25°. Other studies examining co-located talkers reported smaller but significant improvements in speech recognition thresholds (SRT) when masker head orientation angles increased from 45° to 60° (Monson et al., 2019; Trine and Monson, 2020). Children also showed improved SRTs with increasing masker head orientation angle from 45° to 60° (Flaherty et al., 2021). While these previous studies have demonstrated effects of talker head orientation at select angles, no systematic investigation has examined how changes across a full range of orientations affect speech-in-speech recognition. The present study examines the THOR benefit as a function of masker talker head angle for female and male speech.

Because extended high frequencies (EHFs; >8 kHz) exhibit the most directional radiation (Trine et al 2025; Monson et al 2012a), some THOR information is conveyed by EHF cues. Loss of EHF cues decreases talker head orientation discrimination performance by up to 34% (Moriarty et al 2024; Monson et al 2019). EHFs also play a role in other aspects of speech processing. Access to EHFs improves listeners’ speech localization ability in the vertical plane, including front/back discrimination (Best et al., 2005) and improves speech-in-noise recognition when the competing noise is band-limited to 8 kHz (Monson et al., 2023; Motlagh Zadeh et al., 2019; Polspoel et al., 2022). When masker talkers face away from a listener, their EHF energy decreases, providing more glimpsing opportunities for target EHFs (Figure 1B and 1C). Thus, EHF cues provide a benefit for speech-in-speech recognition with a facing target and non-facing masker talkers (Monson et al., 2019; Trine and Monson, 2020; Ananthanarayana et al., 2024).

Individuals with normal audiometric thresholds in the conventional range (0.25 – 8 kHz) but elevated audiometric thresholds at EHFs report and/or demonstrate difficulty with speech recognition in noise (Ananthanarayana et al., 2024; Badri et al., 2011; Helfer et al., 2024; Mishra et al., 2022; Monson et al., 2019; Motlagh Zadeh et al., 2021, 2025; Trine and Monson, 2020). This difficulty may reflect reduced access to the phonetic information conveyed by EHFs, challenges in segregating competing talkers (Monson et al., 2023), or underlying auditory dysfunction such as cochlear synaptopathy (Hunter et al., 2020). In an exploratory analysis, Trine and Monson (2020) reported a correlation between thresholds at 12.5 and 16 kHz and average SRTs for 45° and 60° non-facing maskers. An association between SRTs and 16-kHz thresholds was also reported by Braza et al. (2022) for 60° non-facing masker conditions and by Ananthanarayana et al. (2024) for both facing and non-facing masker conditions. Other studies have reported associations between average EHF thresholds and speech-in-speech performance for traditional facing conditions (Helfer et al., 2024; Mishra et al., 2022;), even though the Speech-in-Noise (SPIN; Bilger et al., 1984) stimuli used by Helfer et al. (2024) had spectral degradations at extended high frequencies (Monson and Buss, 2022) and those used by Mishra et al. (2022) were bandlimited to 11 kHz. However, other studies have found no significant correlation between EHF thresholds and SRTs in facing conditions (Badri et al., 2011; Monson et al., 2023; Smith et al., 2019), although the QuickSIN stimuli used by Smith et al. (2019) had spectral degradations at extended high frequencies (Monson and Buss, 2022) and those used by Badri et al. (2011) were bandlimited to 8 kHz.

Most studies examining THOR or EHF benefit have used female speech (Ananthanarayana et al., 2024; Braza et al., 2022; Flaherty et al., 2021; Monson et al., 2019; Trine and Monson, 2020), whereas relatively few have used male speech (Polspoel et al., 2022; Strelcyk et al., 2014). Long-term average speech spectrum analyses indicate that female speech tends to have higher spectral levels at EHFs than male speech (Delaram et al., 2024; Monson et al., 2012b). However, the perceptual implications of this spectral difference for speech-in-speech recognition are unclear, particularly when target and masker talkers have different head orientations.

Although prior studies have explored THOR benefits and related factors such as availability of EHF cues, EHF thresholds, and talker gender, these investigations have generally included only a limited range of head orientations. In the present study, we systematically evaluated the influence of THOR cues on speech perception for eight masker head orientations ranging from 0° to 180°. We also evaluated the effects of EHF audibility by examining the relationship between performance and 16-kHz audiometric thresholds, and by comparing performance for stimuli that were either full-band or low-pass filtered at 8 kHz in two separate experiments. Experiment 1 used female speech, whereas Experiment 2 used male speech. Based on prior findings, we hypothesized that increasing the head orientation angle, preserving EHF energy, and better 16-kHz thresholds would be associated with better speech recognition performance.

## II. METHODS

### A. Subject

Eighty-three participants were recruited: forty-four participants for the female speech experiment (37 female, 6 male, 1 other; age range = 18-34 years, mean = 21.3) and thirty-nine for the male speech experiment (33 female, 5 male, 1 other; age range = 18-30 years, mean = 21.1). For the female speech experiment, inclusion criteria required participants to be native speakers of American English and have typical hearing indicated by pure-tone air conduction thresholds ≤ 20 dB HL in at least one ear at audiometric frequencies (0.5, 1, 2, 3, 4, 6, 8 kHz) and extended high frequencies (9, 10, 11.2, 12.5, 14, and 16 kHz). Inclusion criteria were the same for the male speech experiment but thresholds at 250 Hz were also considered. Pure-tone audiometric thresholds were obtained using a GSI AudioStar Pro^TM^ audiometer and Radioear DD450 circumaural headphones. Six participants for the female speech experiment and two participants for the male speech experiment had one or two thresholds of 25 or 30 dB HL in one ear. Figure 2 shows boxplots of audiometric thresholds for participants in both experiments. All experimental procedures were approved by the Institutional Review Board at the University of Illinois Urbana-Champaign.

**Figure 2.**
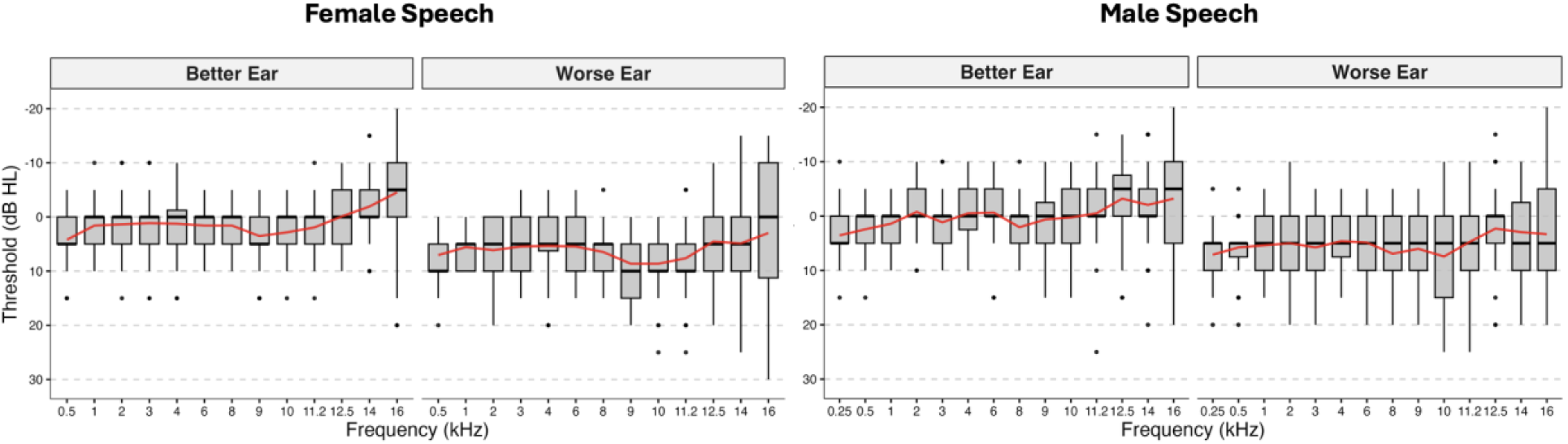
Audiometric thresholds for the better ear and worse ear across test frequencies for the female speech experiment (left) and the male speech experiment (right) participants. Whiskers extend to the most extreme values within 1.5 times interquartile range and points beyond the whiskers indicate outliers. Red lines connect the mean thresholds for each frequency.

### B. Stimuli and conditions

The speech materials were drawn from a publicly available corpus of high-fidelity multi-directional recordings of native speakers of American English (Miller et al., 2024; Monson et al., 2022). The recordings were obtained using seventeen free-field condenser microphones with a flat frequency response up to 20 kHz, arranged in a semicircle with a 1-meter radius. The microphones were positioned from 0° (directly in front of the mouth) to 180° (directly behind the talker’s head) with 11.25° angular separations around the left side of the talker. The talker’s mouth and the microphones were aligned at the same height, with a 1-meter distance between them. Recordings were made at a 48-kHz sampling rate with 24-bit resolution. Recordings from different angles were used to simulate distinct head orientations. This approach preserves natural, dynamic directivity patterns of speech, in contrast to other simulated head orientation methods of filtering on-axis recordings or changing the orientation of a loudspeaker presenting on-axis recordings. However, this approach does not preserve binaural cues related to talker head orientation (see Moriarty et al., 2024). Stimuli were generated based on recordings form three female talkers (target talker 2269 and masker talkers 2258 and 2263 from the corpus) for the female speech experiment and three male talkers (target talker 2273 and masker talkers 2264 and 2281 from the corpus) for the male speech experiment.

Target stimuli were 0° recordings of digits from 0 to 9. Digit triplets were constructed by randomly selecting, level-normalizing, and concatenating the individual digit recordings. Hanning ramps (50 ms) were applied at the boundaries between digits to prevent artificial clicks. There were 150-ms gaps between each digit, and 750-ms of silence at the beginning and end of the triplet. The average duration of a single digit was about 0.72 s (range: 0.53 – 0.92 s), and the digit triplets’ duration was varied based on the duration of selected digits (range with added silences: 3.4 s – 4.5 s). Homogenization of digit recognition difficulty using level adjustment was not conducted because homogenization factors calculated using this approach would depend on masker spectral characteristics. Because masker spectra were changing drastically for the conditions tested here, we elected to equate digits in sound level rather than intelligibility level.

The two-talker masker consisted of segments from unscripted narrative speech (∼2.5 minutes). Recordings from microphones located at 0°, 22.5°, 33.75°, 45°, 67.5°, 90°, 135° and 180° were used to simulate different head orientation angles. For each trial, a segment from each narrative recording was randomly selected, matched in duration to the digit triplet (including leading and lagging silence), and normalized to the same level as the single digits. These segments were mixed with the target speech, and the resulting stimulus was normalized to 72 dB SPL. Non-facing masker recordings were normalized to the same overall level as the facing masker to eliminate the influence of overall-level cues introduced by head orientation, which could reduce masker level as much as 6 dB at 180° (Monson et al 2012), and to isolate spectral mismatch associated with directivity. Figure 3 displays the long-term average speech spectra of the target (digits 0 to 9) and masker recordings for both experiments, plotted on an equivalent rectangular bandwidth (ERB) scale.

**Figure 3.**
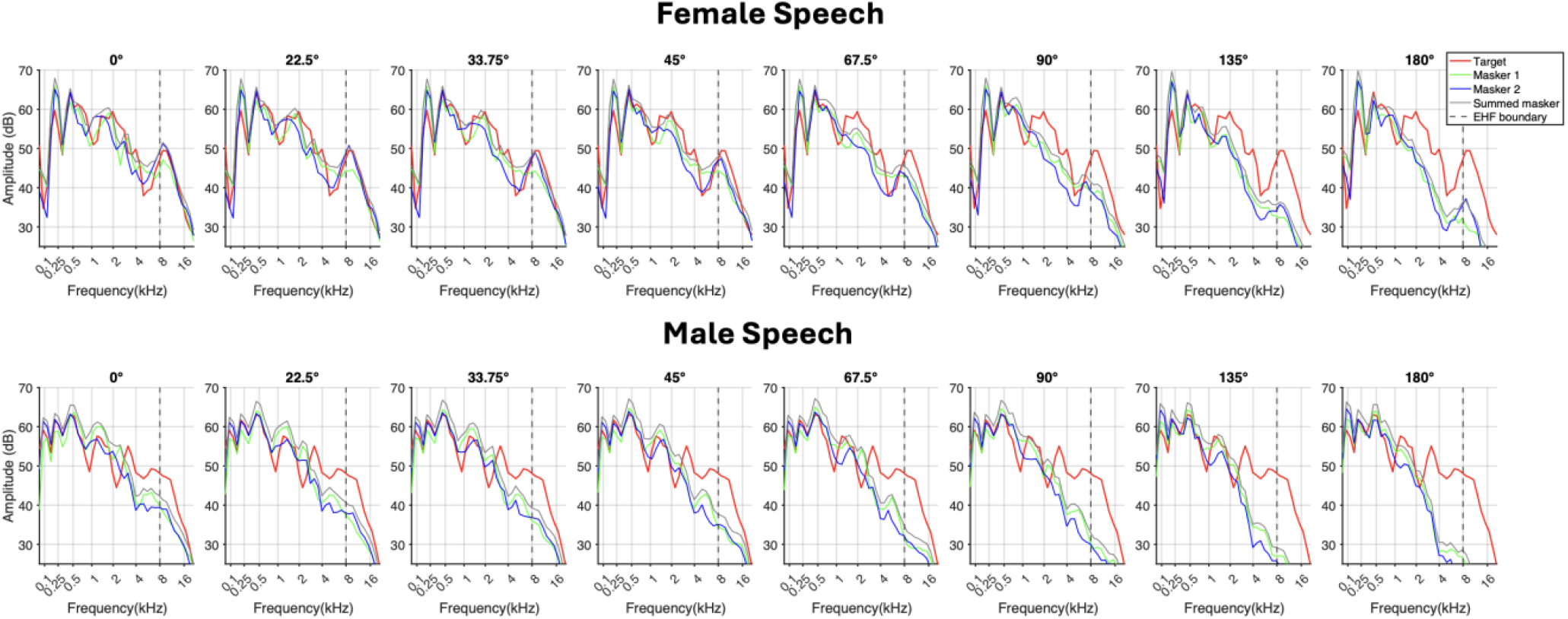
ERB-scaled long-term average speech spectra of recordings used to create target and masker stimuli for female speech (upper panel) and male speech (lower panel). Each recording was normalized to 72 dB SPL and the summed masker stream was 75 dB SPL. Color reflects the talker, as defined in the legend, and a dashed vertical line indicates the 8-kHz boundary between standard and extended high-frequency content.

Eight head orientations and two filtering conditions were used, resulting in a total of 16 experimental conditions. Stimuli were low-pass filtered using a 32-pole Butterworth filter at 8 kHz for low-pass filtered conditions and a 16-pole Butterworth filter at 20 kHz for full-band conditions. Filtering was performed after level normalization to preserve spectral features below 8 kHz. Because energy in the EHF range contributes minimally to the overall stimulus level, level differences introduced by filtering were negligible (< 0.1 dB).

Loudspeaker calibration was conducted using speech-shaped noise generated from the 0° recordings of the masker talkers. For each experiment, a sound level meter was positioned 1 meter from the KRK Rokit 8 loudspeaker to measure the output level. The calibration factor was determined as the difference between the intended presentation level and the level measured by the sound level meter. This calibration factor was subsequently applied to stimuli across all experimental conditions.

### C. Procedure

Following audiometry, participants were seated in a sound-treated booth facing a loudspeaker positioned 1 m in front of them. They were instructed to look at the loudspeaker and limit their movement during stimulus presentation. Participants were instructed to identify the three digits spoken by the target talker while ignoring the maskers. The experiment was implemented in MATLAB (“MATLAB,” R2023a) using custom scripts. A graphical interface displayed the entered digits, mirrored on an iPad positioned in front of the participant using SplashTop software (“Splashtop,” 2023). Participants responded via a wireless keyboard. Each block of trials began with an initial signal-to-noise ratio (SNR) of +4 dB, and SNR changed across the trials in a two-down, one-up adaptive track (Levitt, 1971). The SNR used for each trial was defined as the level of a single target digit relative to one masker; thus the SNR relative to the sum of the two masker stream is 3 dB lower than that reported below. The combined stimulus presentation level was always set to 72 dB SPL to avoid excessively high target SPLs that can trigger acoustic reflexes (Wilson, 1979), while also presenting stimuli at a sufficiently high level to support EHF audibility. Note that this procedure removes natural overall level reductions associated with a head turn, forcing listeners to rely only on spectral distribution THOR cues. The initial step size was 8 dB, reduced to 4 dB after the first reversal, and to 2 dB after the second reversal. Each block ended after total eight reversals, and the block SRT was calculated as the mean of the last six reversals, excluding the first two. The experiment consisted of 16 blocks and started after a practice run. Experimental conditions were presented in random order.

### D. Statistical analysis

All data analyses were conducted using RStudio software (Posit team, 2023). Repeated measures analysis of covariance (ANCOVA; Kassambara, 2026) models were used to examine the effect of the masker head orientation, filtering as within the subjects variable, mean 16-kHz hearing thresholds as covariate, and their interactions on the SRT. Separate and combined models were used for results obtained using female speech and male speech. The assumption of sphericity was assessed using Mauchly’s test. When the assumption was violated, Greenhouse–Geisser corrections were applied to the degrees of freedom and corresponding p values (Greenhouse and Geisser, 1959). A nonlinear mixed-effects model (nlme; Pinheiro et al., 2026) was also used to extrapolate expected performance at head orientation angles not tested here using the combined datasets.

## III. RESULTS

We evaluated speech recognition across eight head-orientation angles to determine how the THOR benefit varies with increasing angle. THOR benefit at each head-orientation angle was defined as the difference between SRTs for that angle and the facing (0°) condition. For both female and male speech, mean SRTs decreased and THOR benefit increased with increasing masker head orientation angle (Figures 4 and 5). For full-band speech, masker head orientations of 67.5° or higher resulted in significant speech recognition performance improvement relative to the facing condition (Figure 5).

**Figure 4.**
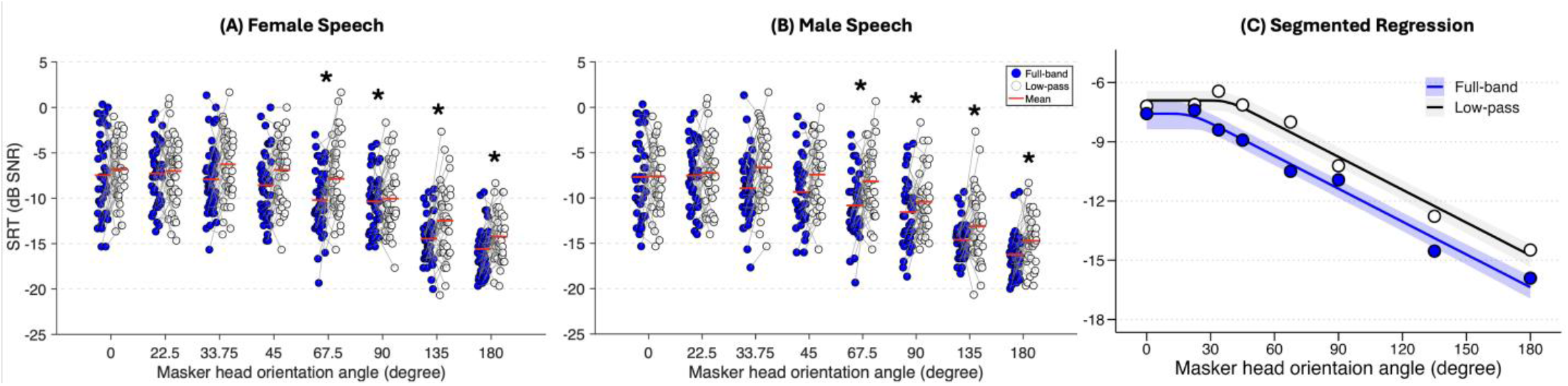
Mean SRT (red dashes) and individual subject data (circles) are plotted for each talker head orientation angle as a categorical variable for female speech (A) and male speech (B) and as a continuous variable as a function of SRT (C). Symbol fill indicates data obtained with stimuli that were full-band (blue) and low-pass filtered at 8 kHz (white). Gray lines connect data points from individual participants across filtering conditions. Asterisks show significant SRT change relative to 0°. Solid lines in right panel indicate predictions from the nonlinear mixed-effects broken-stick model, and shaded regions denote 95% confidence intervals.

**Figure 5.**
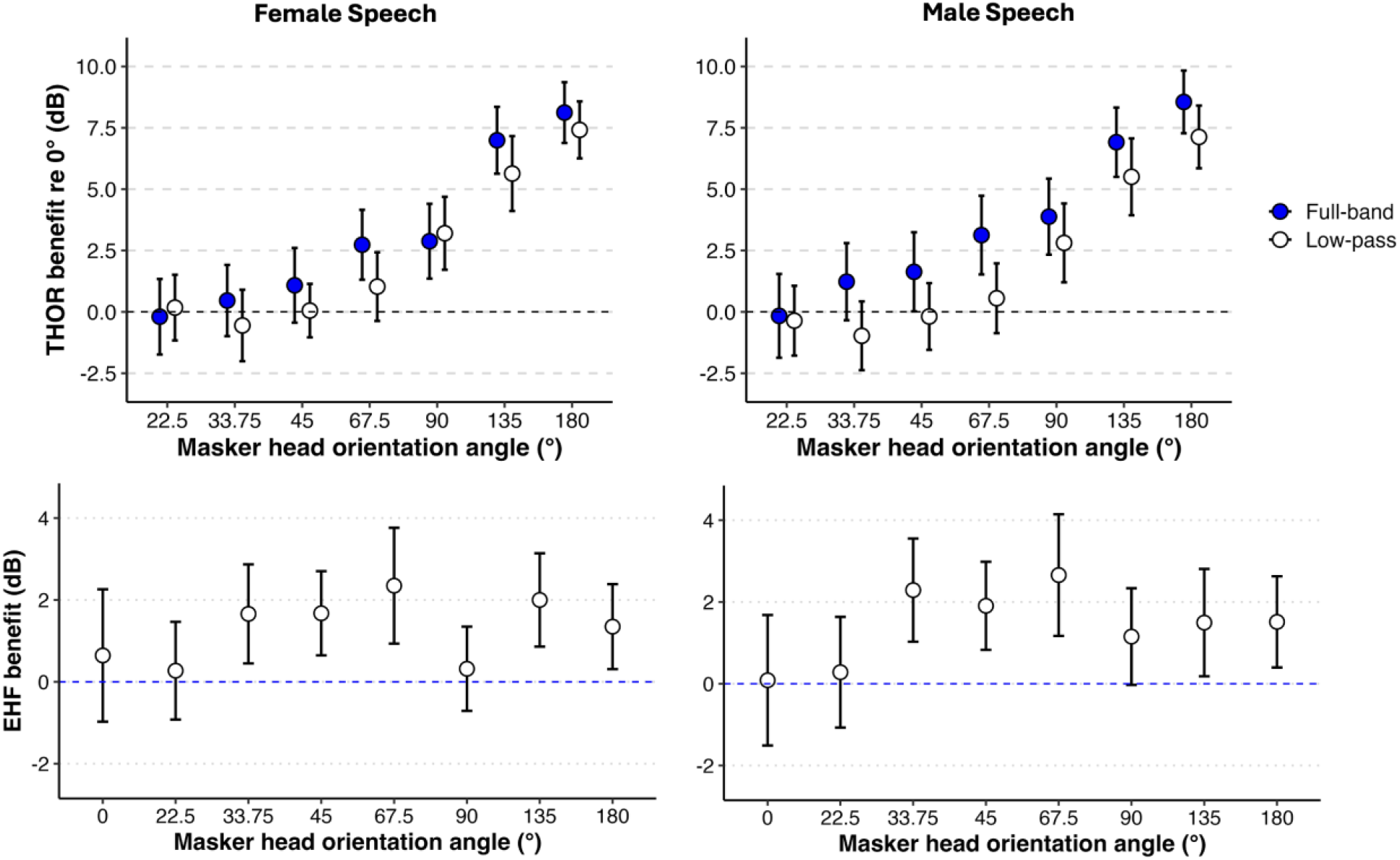
THOR benefit (upper panels) and EHF benefit (lower panels) relative to 0° for female (left) and male (right) speech. Error bars show 95% confidence intervals.

Figure 5 shows EHF benefit at each head-orientation angle, defined as the difference in SRT between the full-band and low-pass conditions. For female speech, mean EHF benefits of ∼2 dB were observed at 33.75°, 45°, 67.5°, and 135°, with a smaller benefit (∼1.4 dB) at 180°. A similar pattern was observed for male speech, with the largest EHF benefits at 33.75°, 45°, 67.5°, followed by 135°, 180°, and 90°. No EHF benefit was observed at 0° or 22.5° in either experiment. It is worth noting that the EHF benefit was highly variable across the participants.

Separate ANCOVAs were conducted for each experiment to assess the effects of head orientation angle, filtering, mean 16-kHz thresholds, and their interactions on SRTs. In both experiments, significant main effects of angle and filtering were observed, while mean 16-kHz thresholds and interaction terms were not significant (Table I). To test for an effect of talker gender, and to increase power to detect potential interactions, both datasets were combined into a single ANCOVA that included talker gender as a between-subjects factor. The model revealed significant effects of angle and filtering, with a significant interaction between angle and filtering (Table II). These results indicate that performance was better for larger masker head angles and for full-band speech, with the interaction indicating THOR and EHF cues influenced one another.

**Table 1.** Results of the ANCOVA models for each experiment. Angle reflects the masker head orientation angle, filtering reflects low-pass filtering at 8 kHz, and PT16kHz is the mean 16-kHz pure-tone threshold for the two ears and is defined as a covariate.

| **Female Speech** |  |  |  |
| --- | --- | --- | --- |
| *Predictors* | *df* | *F* | *p* |
| PT16kHz | 1, 42 | 1.89 | 0.18 |
| Angle | 6, 253.5 | 86.12 | < .001 |
| Filtering | 1, 42 | 45.4 | < .001 |
| PT16kHz × Angle | 6, 253.5 | 0.43 | 0.86 |
| PT16kHz × Filtering | 1, 42 | 2.68 | 0.11 |
| Angle × Filtering | 5.8, 243 | 1.6 | 0.15 |
| PT16kHz × Angle × Filtering | 5.8, 243 | 0.4 | 0.87 |

| **Male Speech** |  |  |  |
| --- | --- | --- | --- |
| *Predictors* | *df* | *F* | *p* |
| PT16kHz | 1, 37 | 1.76 | 0.19 |
| Angle | 5.8, 214.1 | 79.6 | < .001 |
| Filtering | 1, 37 | 52.66 | < .001 |
| PT16kHz × Angle | 5.8, 214.1 | 1.52 | 0.17 |
| PT16kHz × Filtering | 1, 37 | < 0.01 | 0.96 |
| Angle × Filtering | 6, 220.4 | 1.88 | 0.09 |
| PT16kHz × Angle × Filtering | 6, 220.4 | 1.18 | 0.32 |

**Table 2.** Results of the ANCOVA model using combined datasets from both experiments. Angle reflects the masker head orientation angle, filtering reflects low-pass filtering at 8 kHz, gender reflects talker gender, and PT16kHz is the mean 16-kHz pure-tone threshold for the two ears. Gender was included as a between-subject variable, and PT16kHz was a covariate.

| **Combined Model** |  |  |  |
| --- | --- | --- | --- |
| *Predictors* | *df* | *F* | *p* |
| PT16kHz | 1, 80.0 | < 0.01 | 0.99 |
| Gender | 1, 80.0 | 1.48 | 0.23 |
| Angle | 6, 483.7 | 164.29 | < .**001** |
| Filtering | 1, 80.0 | 96.21 | < .**001** |
| PT16kHz × Angle | 6, 483.7 | 0.75 | 0.61 |
| Gender × Angle | 6, 483.7 | 0.13 | 0.99 |
| PT16kHz × Filtering | 1, 80.0 | 1.25 | 0.27 |
| Gender × Filtering | 1, 80.0 | 0.3 | 0.58 |
| Angle × Filtering | 6, 479.0 | 3.27 | < .**01** |
| PT16kHz × Angle × Filtering | 6, 479.0 | 0.74 | 0.62 |
| Gender × Angle × Filtering | 6, 479.0 | 0.29 | 0.94 |

Post-hoc pairwise comparisons were conducted using paired t-tests with Benjamini-Hochberg correction for multiple comparisons (Benjamini and Hochberg, 1995) to investigate SRT differences across head-orientation angles (Table 3). In both experiments, a significant (*p* < 0.05) THOR benefit (compared to the facing condition) first emerged at 67.5° and persisted at larger angles (see Figure 5). Additionally, SRTs for 67.5° and beyond were significantly better than SRTs for 22.5°, 33.75°, and 45°. Pairwise comparisons between all angles ≥67.5° were significant, indicating that each increase in angle beyond 67.5° led to significantly improved speech recognition performance.

**Table 3.**
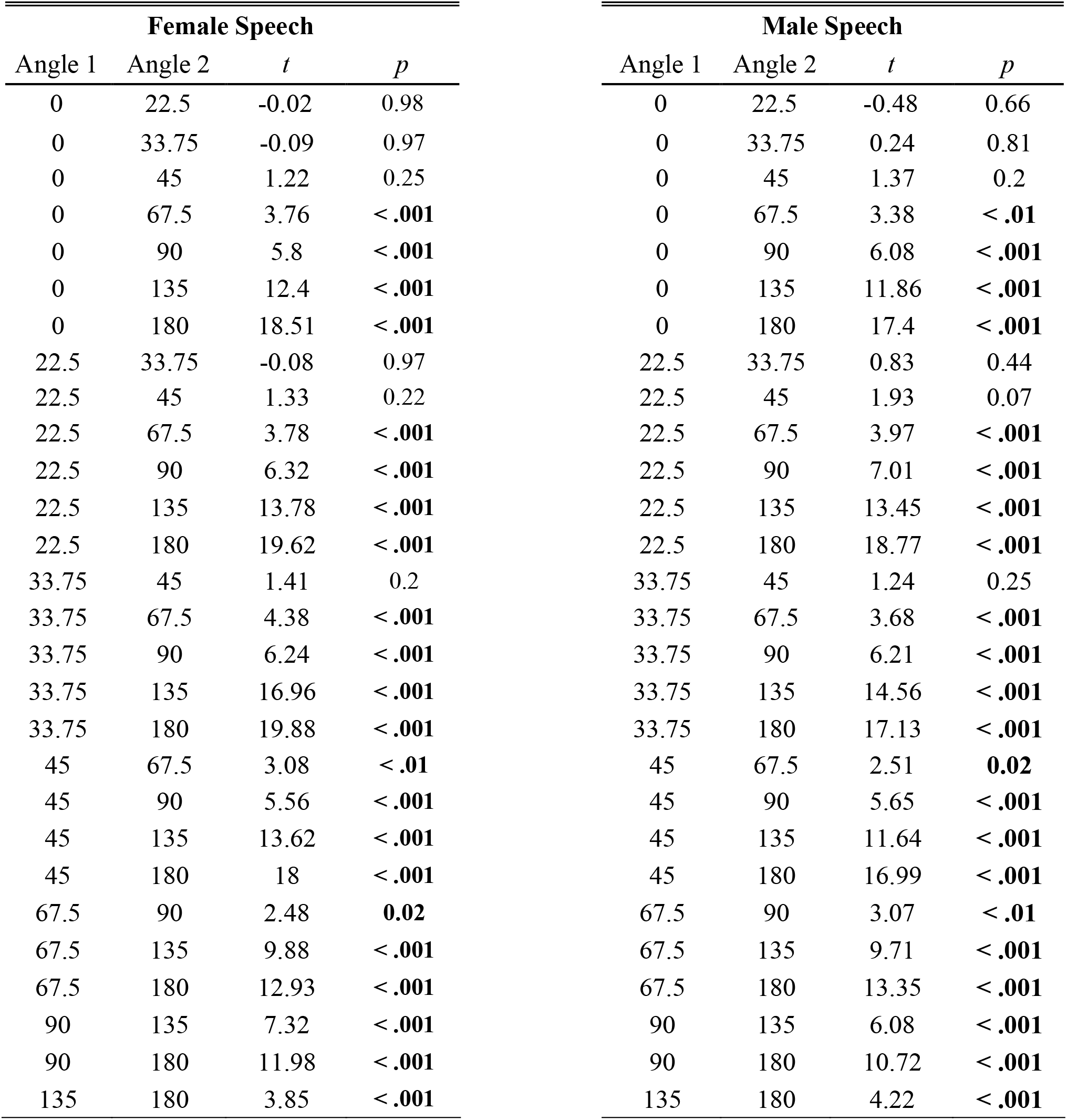
Results of post-hoc pairwise comparisons of different masker head orientation angles for female and male speech. P-values were adjusted for multiple comparisons using the Benjamini- Hochberg method.

| **Female Speech** |  |  |  |
| --- | --- | --- | --- |
| Angle 1 | Angle 2 | *t* | *p* |
| 0 | 22.5 | -0.02 | 0.98 |

| **Male Speech** |  |  |  |
| --- | --- | --- | --- |
| Angle 1 | Angle 2 | *t* | *p* |
| 0 | 22.5 | -0.48 | 0.66 |

Table 3. Results of post-hoc pairwise comparisons of different masker head orientation angles for 265 female and male speech. P-values were adjusted for multiple comparisons using the Benjamini- 266 Hochberg method.
|  |  |  |  |  |  |  |  |
| --- | --- | --- | --- | --- | --- | --- | --- |
| 0 | 33.75 | -0.09 | 0.97 | 0 | 33.75 | 0.24 | 0.81 |
| 0 | 45 | 1.22 | 0.25 | 0 | 45 | 1.37 | 0.2 |
| 0 | 67.5 | 3.76 | < .001 | 0 | 67.5 | 3.38 | < .01 |
| 0 | 90 | 5.8 | < .001 | 0 | 90 | 6.08 | < .001 |
| 0 | 135 | 12.4 | < .001 | 0 | 135 | 11.86 | < .001 |
| 0 | 180 | 18.51 | < .001 | 0 | 180 | 17.4 | < .001 |
| 22.5 | 33.75 | -0.08 | 0.97 | 22.5 | 33.75 | 0.83 | 0.44 |
| 22.5 | 45 | 1.33 | 0.22 | 22.5 | 45 | 1.93 | 0.07 |
| 22.5 | 67.5 | 3.78 | < .001 | 22.5 | 67.5 | 3.97 | < .001 |
| 22.5 | 90 | 6.32 | < .001 | 22.5 | 90 | 7.01 | < .001 |
| 22.5 | 135 | 13.78 | < .001 | 22.5 | 135 | 13.45 | < .001 |
| 22.5 | 180 | 19.62 | < .001 | 22.5 | 180 | 18.77 | < .001 |
| 33.75 | 45 | 1.41 | 0.2 | 33.75 | 45 | 1.24 | 0.25 |
| 33.75 | 67.5 | 4.38 | < .001 | 33.75 | 67.5 | 3.68 | < .001 |
| 33.75 | 90 | 6.24 | < .001 | 33.75 | 90 | 6.21 | < .001 |
| 33.75 | 135 | 16.96 | < .001 | 33.75 | 135 | 14.56 | < .001 |
| 33.75 | 180 | 19.88 | < .001 | 33.75 | 180 | 17.13 | < .001 |
| 45 | 67.5 | 3.08 | < .01 | 45 | 67.5 | 2.51 | 0.02 |
| 45 | 90 | 5.56 | < .001 | 45 | 90 | 5.65 | < .001 |
| 45 | 135 | 13.62 | < .001 | 45 | 135 | 11.64 | < .001 |
| 45 | 180 | 18 | < .001 | 45 | 180 | 16.99 | < .001 |
| 67.5 | 90 | 2.48 | 0.02 | 67.5 | 90 | 3.07 | < .01 |
| 67.5 | 135 | 9.88 | < .001 | 67.5 | 135 | 9.71 | < .001 |
| 67.5 | 180 | 12.93 | < .001 | 67.5 | 180 | 13.35 | < .001 |
| 90 | 135 | 7.32 | < .001 | 90 | 135 | 6.08 | < .001 |
| 90 | 180 | 11.98 | < .001 | 90 | 180 | 10.72 | < .001 |
| 135 | 180 | 3.85 | < .001 | 135 | 180 | 4.22 | < .001 |

Whereas ANCOVA models represented head orientation categorically, the present data were also evaluated as a funciton of angle represented numerically. We modeled speech recognition performance for each experiment using nonlinear mixed-effects broken-stick models that were fit to SRTs with masker angle and EHF condition as predictors, with random intercepts for participants (Figure 4C). A broken stick function was selected based on visual inspection of the data, to capture the similarity of SRTs at 0° and 22.5°. Free parameters in this model were the intercept, the breakpoint, and the slope as a function of angle above the breakpoints. Each of these three parameters was allowed to vary by talker gender and filter conditions. Talker gender and filter condition were both sum coded (-0.5 male and 0.5 female; -0.5 low-pass and 0.5 full-band). Results indicated significant effects of intercept (Coef = -7.26, t = -26.50, p <.001), breakpoint (Coef = 30, t = 6.93, p < .001) and slope (Coef = -0.06, t = -29.46, p < .001). Slope did not differ significantly by talker or filter condition, and there was no evidence of an effect of talker gender for either breakpoint or intercept (p ≥ .437). There was a significant effect of filter condition on breakpoint (p = .026), indicating that the improvement in SRT began at a smaller masker angle in the full-band condition than in the low-pass condition. Specifically, the mean estimated breakpoint was 20.5° for the full-band condition and 39.9° for the low-pass condition. There was also a non-significant trend for an effect of filter condition on intercept (p = .079), with predicted values of -7.6 dB for the full-band condition and -6.9 dB for the low-pass condition.

This model captures some of the salient trends in the data, but the overall quality of the model fit is modest, accounting for 42% of variance. Some of the unfitted variance is likely related to measurement noise. For example, there was a 15.7 dB range for SRTs obtained for the 0° masker in the female full-band condition, but a correlation of only r = 0.18 between SRTs for the 0° and 22.5° conditions, with comparable results for the male talker data. Another possible source of error is related to evidence of deviation from linear reduction in SRT as a funciton of angle. For example, the mean SRT associated with 135° falls below the prediction based on lines fits for both talker genders and filter conditions. Predictions based on these fits should therefore be treated with some caution.

## IV. DISCUSSION

In real-world multi-talker environments, talkers typically face their communication partners; as a result, listeners are rarely confronted with multiple simultaneous streams of speech from talkers who are all facing them. In this study, we examined the effect of masker head orientations on speech recognition performance using female and male speech. We also evaluated the contribution of EHFs and 16-kHz audiometric thresholds. For full-band speech, SRTs improved by 2 to 8 dB when masker heads were oriented between 67.5° and 180°, relative to the 0° facing condition. Performance at 67.5° was significantly better than at all smaller angles but poorer than at larger angles, suggesting that the THOR benefit emerges between 45° and 67.5°, with additional improvements as orientation increases beyond this range. Access to EHFs yielded a relatively small but consistent performance enhancement of approximately 1–2 dB for both male and female speech. The angle ’ filtering interaction indicated that the EHF benefit was dependent on head angle and that the THOR benefit was larger for full-band speech than for 8-kHz bandlimited speech (see Figure 5).

### THOR benefit

THOR benefits observed at orientations greater than 45° are likely attributable to the increased spectral mismatches between the target and masker speech signals at these angles. These spectral mismatches can facilitate recognition of the target talker via stream segregation or unmasking of phonetic cues. As masker head orientation increases, target high-frequency energy, particularly at EHFs, becomes progressively unmasked due to its directionality (Monson et al., 2012a; Monson and Ananthanarayana, 2023; Trine et al., 2025; see Figure 3).

The observation that THOR benefit emerged between 45° and 67.5° is consistent with published data. Previous studies reported improved performance when masker head orientation shifted from 0° to 105° (Strelcyk et al, 2014), from 0° to 60° (Braza et al. 2022), and from 0° to 56.25° (Ananthanarayana et al. 2024). It is also consistent with findings of Monson et al. (2019), Trine and Monson (2020), and Flaherty et al. (2021) that increasing masker head orientation from 45° to 60° improved speech recognition performance. Collectively, these results suggest that conventional speech-in-speech paradigms using exclusively front-facing talkers may underestimate real-world speech recognition performance, where masker orientations are more variable and rarely aligned directly at the listener.

Our findings highlight differences between two types of spatial benefit: THOR benefit arising from mismatched target and masker head orientations, and spatial release from masking (SRM) arising from mismatched target and masker spatial locations. SRM for a target presented in a two-talker masker emerges at small angles (<15° of separation; Marrone et al., 2008; Srinivasan et al., 2016) and has been reported to plateau at 12-13 dB at approximately 45° of separation, with minimal increases beyond this angle (Marrone et al 2008). In contrast, THOR benefit doesn’t emerge until approximately 45° of head rotation and steadily increases to 8 dB at 180° (Figure 4). Based on these datasets, one might conclude that spatial separation provides a greater speech recognition benefit than mismatched head orientation (12-13 dB vs. 8 dB). However, in the present study, natural overall-level reductions associated with non-facing masker head orientations were not available to listeners due to stimulus level normalization, which scaled all maskers to the same level as the facing (0°) masker. Incorporating natural overall-level reductions associated with non-facing masker orientations would likely increase the THOR benefit (Strelcyk et al., 2014). For example, for the stimuli used in the present study, the 180° masker would have been 5.68 dB lower in level than the 0° masker. Assuming a strong (potentially one-to-one) correspondence between masker sound level and SRT, we speculate that the THOR benefit at 180° could approach or exceed 12-13 dB with this masker level reduction.

### EHFs and EHF Thresholds

The EHF benefit likely reflects the contribution of EHF temporal and spectral cues to phonetic information and/or stream segregation (Trine and Monson, 2020; Monson et al 2023). This benefit was generally observed for masker head orientation angles >22.5°, but was reduced at 90°. Due to greater masking energy at EHFs associated with facing maskers, the lack of EHF benefit for smaller masker head angles was expected, but the reduced EHF benefit at 90° was not expected. It is not clear why a 90° masker head orientation would reduce the benefit of EHF cues, but this pattern was observed for both experiments. One possibility we considered is that 90° head orientation maskers may selectively unmask some lower-frequency spectral regions such that the reliance on EHF cues is reduced. However, evidence to support this possibility is not apparent from the acoustical analysis shown in Figure 3. Another possibility is that some EHFs exhibit increased radiation at 90° for the masker talkers used in this study. However, Figure 3 and directivity analyses previously conducted on these talkers (Trine et al., 2025) do not support this explanation either. This phenomenon warrants further investigation.

There was no significant relationship between 16-kHz thresholds and SRTs in either dataset. This may be because all listeners here were young, normal-hearing listeners with relatively good EHF hearing (mean 16-kHz thresholds ≤20 dB HL). However, this finding stands in contrast with the findings of Trine and Monson (2020) who reported a correlation between 16-kHz thresholds and SRTs using a similar speech recognition paradigm for a similarly young, normal-hearing demographic with good EHF hearing. Braza et al. (2022) and Ananthanarayana et al. (2024) also showed an effect of EHF sensitivity at 16 kHz on SRTs using a similar approach, however those studies included listeners with poor EHF hearing, resulting in a wider range of EHF thresholds. It may be that we did not observe a relationship here because of the limited range of EHF thresholds. By the same token, claims surrounding the EHF and THOR benefits observed here are limited to those with good EHF hearing.

The present study had additional limitations. Maskers in this study were co-located to establish baseline estimates of THOR benefit. Notably, most previous studies of SRM used facing maskers. In the real world, maskers are spatially separated and non-facing, leading to an interaction influencing the magnitude of THOR benefit and SRM (Braza et al., 2022) and this phenomenon warrants further investigation. We used only one female target talker and one male target talker who had similar EHF spectral levels. Consequently, these findings (including the lack of a gender effect) may not be generalizable to other female and male talkers, especially considering that male talkers tend to have lower EHF levels than female talkers, on average (Delaram et al. 2024). Whereas the stimuli captured spectral effects of head orientation, they did not include differences in speech level, effects of reverberation and distance, or the natural binaural cues associated with non-facing orientations of directional sources. Future research should explore the combined effect of these additional factors on speech recognition. Such work would help inform the design of real-world listening experiments and contribute to more accurate models of auditory scene analysis.

## V. CONCLUSION

This study systematically examined the effects of masker head orientation, EHF cues, and 16- kHz audiometric thresholds on speech recognition in multi-talker conditions using both female and male speech. THOR benefits emerged at a masker head orientation > 45°, with progressively greater improvements at larger angles. Access to EHFs provided a small but consistent speech recognition benefit for a facing target talker and masker head orientation angles >22.5°.

## ACKNOWLEDGMENTS

We thank our study participants. This project was supported by the National Institutes of Health Grant No. R01-DC019745 (B.B.M.).

## AUTHOR DECLARATIONS

### Conflict of Interest

The authors declare no conflict of interest.

### Ethics Approval

Informed consent was obtained from all participants, and all data collection procedures were approved by the Institutional Review Board at the University of Illinois at Urbana-Champaign (Protocol 24-1627).

## DATA AVAILABILITY

The data that support the findings of this study are available upon request.

## Notes

### Competing Interest Statement

The authors have declared no competing interest.

